# Tissue Specific Knockout of the Cardiolipin Transacylase Enzyme TAFAZZIN in Both Liver and Pancreatic Beta Cells Protects Mice From Diet-Induced Obesity

**DOI:** 10.1101/2022.04.19.488810

**Authors:** Laura K. Cole, Vernon W. Dolinsky, Grant M. Hatch

**Affiliations:** Department of Pharmacology & Therapeutics, University of Manitoba, Children’s Hospital Research Institute of Manitoba, Winnipeg, Manitoba, Canada, R3E3P4

## Abstract

Mutations in the TAFAZZIN gene result in the X-linked genetic disease Barth Syndrome. The protein product tafazzin is a transacylase enzyme that remodels the phospholipid cardiolipin with fatty acids. Some Barth syndrome boys exhibit a lean phenotype. In addition, whole body knockdown of tafazzin in mice protects them from diet-induced obesity through increased hepatic fatty acid oxidation and reduced basal insulin secretion. We thus hypothesized that tafazzin deficiency in both the liver and beta cells of the pancreas contribute to this lean phenotype. Through a Cre-Lox approach we generated control, liver-specific, pancreatic beta cell-specific and liver and pancreatic beta cell-specific double knockout male mice. The animals were fed a high fat diet for 8 weeks and body weight, liver weight and fat pad weights determined. Liver-specific or pancreatic beta cell-specific male tafazzin knockout mice accumulated weight gain (≍40% increase in body weight) at the same rate as control animals. In contrast, the liver- and beta cell-specific double tafazzin knockout mice exhibited a reduced rate of weight gain by 8 weeks (≍26% increase in body weight) compared to control or the single tafazzin knockout animals. In addition, at 8 weeks the double tafazzin knockout mice exhibited reduced weight gain in tissues known to accumulate fat including the liver, the gonadal, inguinal and perirenal white adipose tissues and the brown adipose tissue. Thus, liver- and pancreatic beta cell-specific double tafazzin knockout male mice are protected from high fat diet induced weight gain and fat accumulation. These results may partially explain why some Barth Syndrome boys exhibit a lean phenotype.

## Introduction

Diet-induced obesity results in mitochondrial defects that contribute to development of tissue steatosis and metabolic complications^1^. Cardiolipin (CL) is a key mitochondrial lipid that regulates mitochondrial bioenergetics^2^. Alteration in CL is directly linked to development of obesity^1^. Barth Syndrome (BTHS) is a rare X-linked disorder in which the specific defect is a reduction in CL^3^. The loss in CL is caused by mutation in the TAFAZZIN gene which codes for tafazzin (Taz), a protein that synthesizes CL. Interestingly, BTHS boys possess a lean phenotype with mean weight in the 15^th^ percentile and body mass index below the 5^th^ percentile^4^. We previously showed that mice with whole body knockdown of Taz are protected from diet-induced obesity^5^. In addition, pancreatic islets isolated from Taz knockdown mice are protected from obesity-induced beta cell dysfunction^6^. The reduction in CL seen in most tissues of Taz knockdown mice was not mirrored in the liver^1^ nor in pancreatic islets^6^. As a result, Taz knockdown mice exhibited normal hepatic mitochondrial function and, in combination with a reduction in basal insulin secretion from pancreatic islets, contributed to metabolic conditions consistent with a lean phenotype^5,6^. Recently, we demonstrated that treatment of high fat diet fed Taz knockdown mice with resveratrol, a small cardioprotective nutraceutical, further attenuated the aberrant lipid accumulation in their hearts^7^. Collectively, these studies identified a novel role for Taz in regulating susceptibility to diet-induced obesity. However, the mechanism by which Taz knockdown conferred protection to diet-induced obesity mediated liver and beta cell dysfunction was unclear since Taz was reduced in all tissues of these animals.

Based upon our previous observations ^5–7^, we hypothesized that a targeted deletion of Taz in both the liver and pancreatic beta cells in mice would protect them from diet-induced obesity and tissue steatosis. We utilized the Cre/lox system for generating tissue-specific knockout and our results show that liver- and pancreatic beta cell-specific double tafazzin knockout male mice are indeed protected from high fat diet-induced weight gain and fat accumulation.

## Materials and Methods

Embyros from mice containing lox*P*-flanked Taz (flox) were previously generated by using sperm from a Taz flox male (Taz-fl/Y) for *in vitro* fertilization of wild type C57BL/6J oocytes (Dr. Douglas Strathdee, Beaton Institute, Glasgow, UK). The resulting heterozygous female (Taz-fl/+) and hemizygous males (Taz-fl/Y) were bred >9 times to a C57BL/6 background. To specifically delete Taz in the liver, mice homozygous/hemizygous for the floxed Taz allele are mated with B6.Cg-Speer6-ps1^Tg(Alb-cre)21Mgn^/J strain of mice transgenic for cre recombinase under the albumin (*Albumin*)promoter (*AlbuminCre*). The B6.Cg-*Speer6-ps1^Tg(Alb-cre)21Mgn^*/J strain of mice are obtained from Jackson Laboratories. Subsequent breeding took place between female mice heterozygous for the floxed Taz allele (Taz-fl/+, without *AlbuminCre*) with male mice which are wildtype (+/+) as well as male mice which are hemizygous for the floxed Taz allele (Taz-fl/Y) and heterozygous for the *AlbuminCre* transgene. This generated male and female mice which are hemizygous for the floxed Taz allele (Taz-fl/Y) and homozygous for the floxed allele (Taz-fl/Taz-fl) with *AlbuminCre* expression (liver specific knockout of Taz). Littermate controls included both male and female floxed Taz (Taz-fl/Y, Taz-fl/Taz-fl) without *AlbuminCre* expression (flox controls) and wildtype Taz with *AlbuminCre* (cre recombinase controls) (**Figure 1**). The same protocol was used to generate pancreatic beta cell-specific TazKO mice driven under the Ins1 promoter using B6.Cg-*^Tg(Ins1-cre/ERT)1Lphi^*/J strain of mice. In addition, liver- and pancreatic beta cell-specific double knockout mice were generated. 4 Month old male mice were fed a high-fat (23%, w/w) (Envigo Teklad, Rodent diet cat# TD.130261) diet for up to 8 weeks. Body weights were measured weekly and fat accumulating tissues were removed and weighed after 8 weeks. All data are presented as mean+standard error of the mean, n=2-4.

**Figure 1.**
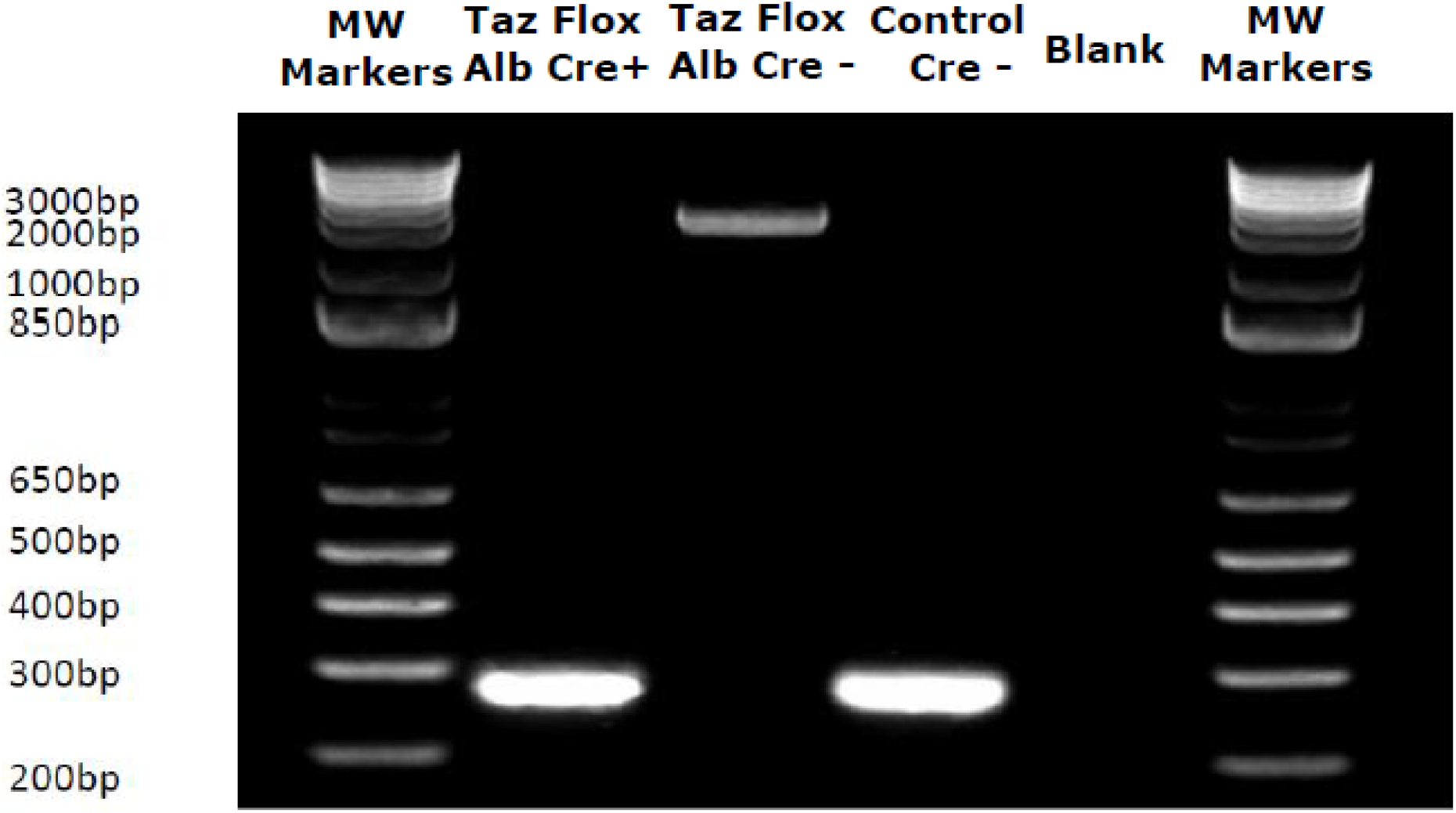
Genotyping of TAFAZZIN knockout mouse liver. Primers for the Taz gene (TazGTKOU1 and TazGTFLoxD1) DNA were isolated from the liver of Floxed mice. Product size for WT Taz 3.1 kb.Product size for KO of Taz 280 bp. A representative gel is depicted.

## Results & Discussion

Control, liver-specific tafazzin knockout, pancreatic beta cell-specific tafazzin knockout, or liver- and pancreatic beta cell-specific tafazzin double knockout male mice were fed a high-fat diet for up to 8 weeks and body weights determined. Liver-specific or pancreatic beta cell-specific male tafazzin knockout mice accumulated weight gain (≍40% increase in body weight) at the same rate as control animals (**Figure 2A**). In contrast, the liver- and beta cell-specific double tafazzin knockout mice exhibited a reduced percentage of weight gain by 8 weeks (≍26% increase in body weight) compared to control or the single tafazzin knockout animals (**Figure 2B**). Thus, liver- and pancreatic beta cell-specific tafazzin double knockout male mice exhibit a reduced weight gain on a high fat diet.

**Figure 2.**
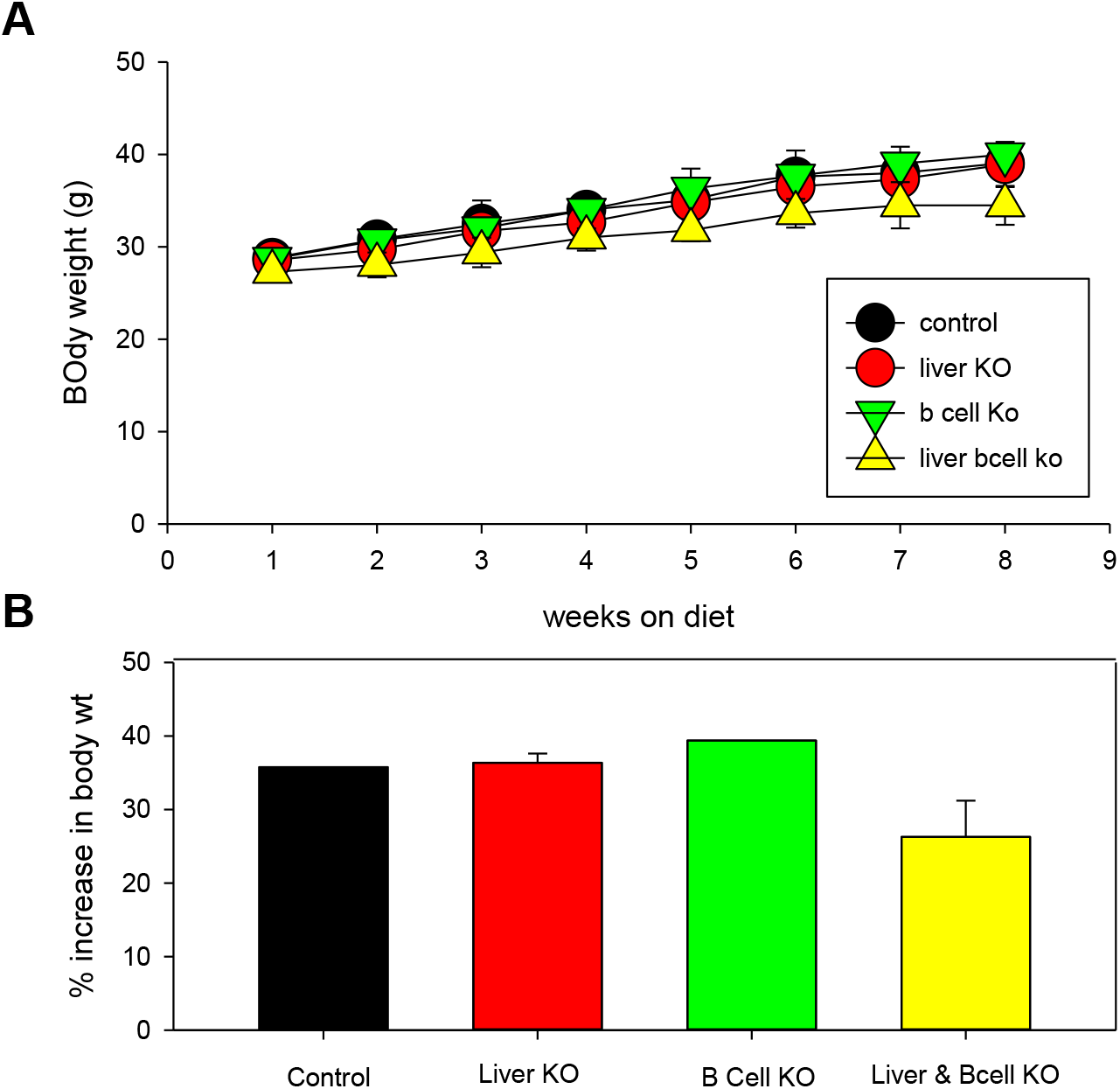
A. Temporal increase in body weight of male mice fed the high fat diet. B. Percent increase in body weight after 8 weeks on the high fat diet (B). Control (Black), liver TazKO (Liver KO) (Red), pancreatic beta cell TazKO (B Cell KO) (Green) and liver- and pancreatic beta cell-specific double TazKO (Liver & Beta KO) (Yellow) male mice. Data represent mean±standard error of the mean, n=2-4.

We examined the overall body weight and weights of the tissues known to accumulate fat including the liver, the gonadal, inguinal and perirenal white adipose tissues (WAT) and the brown adipose tissue (BAT) in these animals. Liver-specific or pancreatic beta cell-specific male tafazzin knockout mice body weight was similar to control animals (**Figure 3A**). In contrast, liver- and beta cell-specific double tafazzin knockout mice exhibited a lowered weight at 8 weeks compared to control or the single tafazzin knockout animals. In addition, at 8 weeks of feeding the high fat diet the double tafazzin knockout mice exhibited reduced weight gain in tissues known to accumulate fat including the liver, the gonadal WAT, inguinal WAT, perirenal WAT and BAT (**Figure 3B-F**).

**Figure 3.**
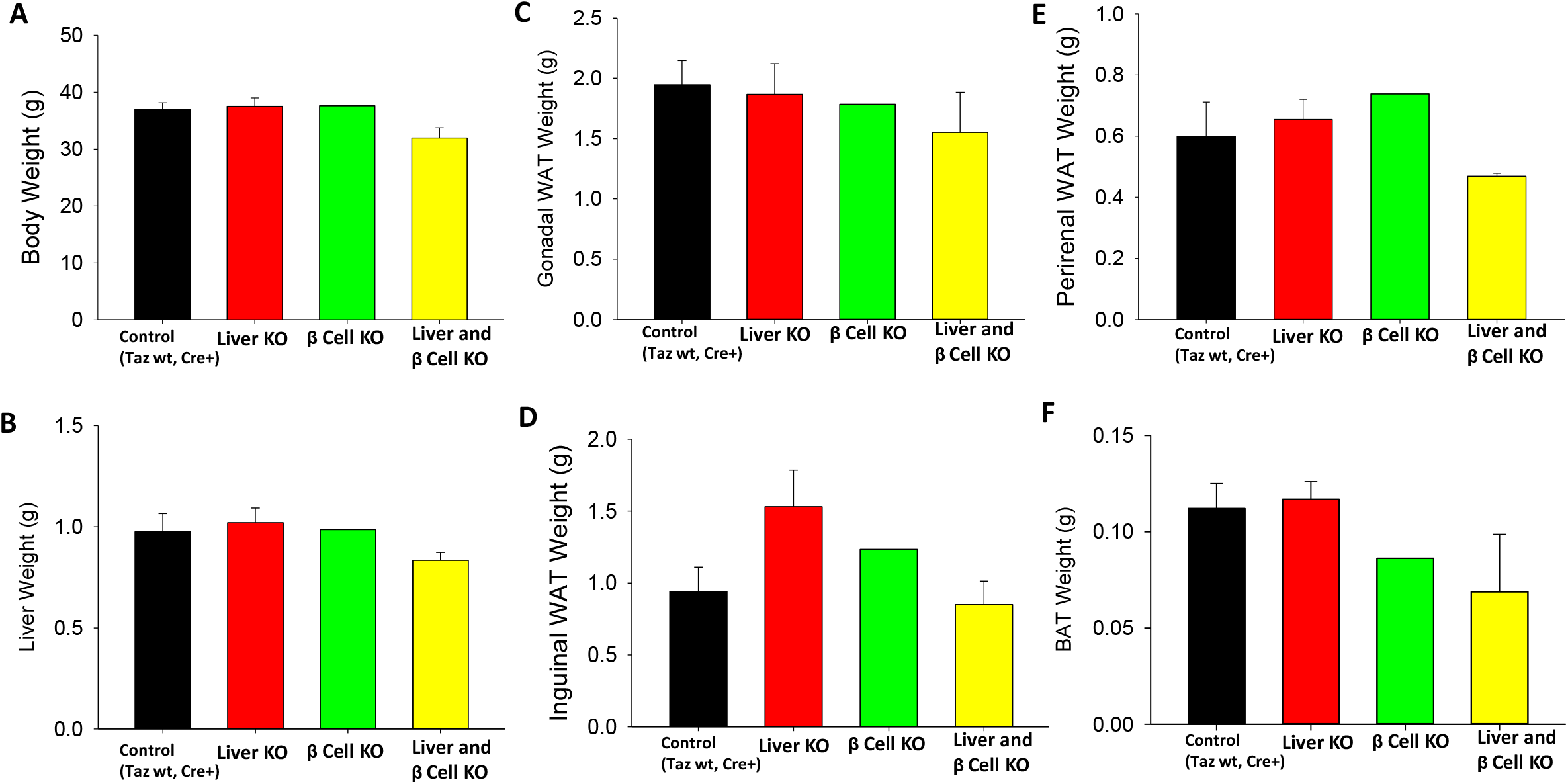
Body weight (A), liver weight (B), gonadal WAT weight (C), inguinal WAT weight (D), perirenal WAT weight (E), and BAT weight in control (Black), liver TazKO (Liver KO) (Red), pancreatic beta cell TazKO (B Cell KO) (Green) and liver- and pancreatic beta cellspecific double TazKO (Liver & Beta KO) (Yellow) male mice fed a high fat diet for 8 weeks. Data represent mean±standard error of the mean, n=2-4.

Liver-specific or pancreatic beta cell-specific male tafazzin knockout mice body exhibited similar total gonadal WAT, inguinal WAT, perirenal WAT and BAT fat pad mass compared to control animals after 8 weeks of high fat diet (**Figure 4**). In contrast, liver- and beta cell-specific double tafazzin knockout mice exhibited lowered gonadal WAT, inguinal WAT, perirenal WAT and BAT fat pad mass compared to control or the single tafazzin knockout animals after 8 weeks of high fat diet. Thus, liver- and pancreatic beta cell-specific double tafazzin knockout male mice are protected from high fat diet induced weight gain and fat accumulation. In conclusion, we have found that liver and pancreatic beta cell Taz double knockout mice are protected from high fat diet-induced weight gain and fat accumulation. These findings complement our previous studies in which TAZ deficiency protects mice from diet-induced obesity^5–7^. Together, our data reveal a novel role for TAZ in regulating diet-induced weight gain and fat accumulation.

**Figure 4.**
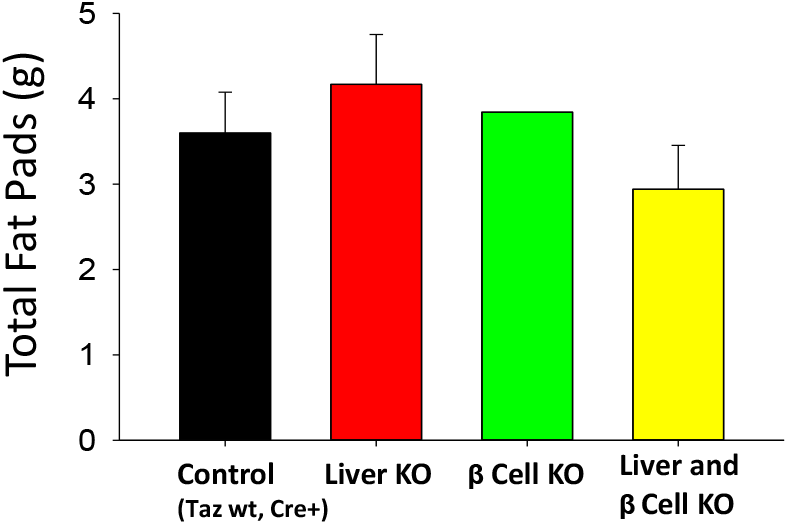
Total fat pad weight in control (Black), liver TazKO (Liver KO) (Red), pancreatic beta cell TazKO (B Cell KO) (Green) and liver- and pancreatic beta cell-specific double TazKO (Liver& Beta KO) (Yellow) male mice fed a high fat diet for 8 weeks. Data represent mean±standard error of the mean, n=2-4.

## Acknowledgements

We thank Dr. Douglas Strathdee, Beaton Institute, Glasgow, UK, for provision of the embyros from mice containing lox*P*-flanked Taz (flox). The technical assistance of Ms. Marilyne Vandel was greatly appreciated. This work was supported by grants from the University of Manitoba Research Grants Program, the Children’s Hospital Research Institute of Manitoba, Heart and Stroke Foundation of Canada, CIHR Impact Training Program. GMH is the CRC in Molecular Cardiolipin Metabolism.

## References

1. Feillet-Coudray C, Fouret G, Casas F, Coudray C. (2014) Impact of high dietary lipid intake and related metabolic disorders on the abundance and acyl composition of the unique mitochondrial phospholipid, cardiolipin. J Bioenerg Biomembr. 46, 447–457.

2. Hatch GM (2004) Cell biology of cardiac mitochondrial phospholipids. Biochem Cell Biol. 82, 99–122.

3. Hauff KD, Hatch GM. (2006) Cardiolipin metabolism and Barth Syndrome. Prog Lipid Res. 45, 91–101.

4. Ferreira C, Thompson R, Vernon H. (2014) Barth Syndrome. In: Adam MP, Ardinger HH, Pagon RA, Wallace SE, Bean LJH, Stephens K, Amemiya A, editors. GeneReviews^®^ [Internet]. Seattle (WA): University of Washington, Seattle; 1993–2019. Oct 9.

5. Cole LK, Mejia EM, Vandel M, Sparagna GC, Claypool SM, Dyck-Chan L, Klein J, Hatch GM. (2016) Impaired Cardiolipin Biosynthesis Prevents Hepatic Steatosis and Diet-Induced Obesity. Diabetes 65, 3289–3300.

6. Cole LK, Agarwal P, Doucette C, Fonseca M, Xiang B, Sparagna GC, Seshadri N, Vandel M, Dolinsky VW, Hatch GM. (2021). Tafazzin Deficiency Reduces Basal Insulin Secretion and Mitochondrial Function in Pancreatic Islets from Male Mice. Endocrinol. 162, 1–15.

7. Cole LK, Meija EM, Sparagna GC, Vandel M, Xiang B, Han X, Dedousis N, Kaufman BA, Dolinsky VW, Hatch GM. (2020) Cardiolipin Deficiency Elevates Susceptibility to a Lipotoxic Hypertrophic Cardiomyopathy. J Mol Cell Cardiol. 144, 24–34.

